# SingleFrag: A deep learning tool for MS/MS fragment and spectral prediction and metabolite annotation

**DOI:** 10.1101/2024.11.04.621329

**Authors:** Maribel Pérez-Ribera, Muhammad Faizan Khan, Roger Giné, Josep M. Badia, Sandra Junza, Oscar Yanes, Marta Sales-Pardo, Roger Guimerà

## Abstract

Metabolite and small molecule identification via MS/MS involves matching experimental spectra with prerecorded spectra of known compounds. This process is hindered by the current lack of comprehensive reference spectral libraries. To address this gap, we need accurate *in silico* fragmentation tools for predicting MS/MS spectra of compounds for which empirical spectra do not exist. Here, we present SingleFrag, a novel deep learning tool that predicts individual fragments separately, rather than attempting to predict the entire fragmentation spectrum at once. Our results demonstrate that SingleFrag surpasses state-of-the-art *in silico* fragmentation tools, providing a powerful method for annotating unknown MS/MS spectra of known compounds. As a proof of concept, we successfully annotate three previously unidentified compounds frequently found in human samples.

## Introduction

Mass spectrometry (MS) is a powerful analytical tool for studying small molecules and metabolites in biological systems. The interpretation of tandem MS (MS/MS) spectra is crucial for metabolite annotation and identification, driving advancements in metabolomics across diverse fields such as precision medicine^1^, biomarker discovery^2^, nutritional sciences^3^, microbiome research^4^, toxicology and environmental testing^5^.

However, practical application of MS/MS for metabolite identification presents significant challenges. Traditional approaches rely on fragmentation libraries^6^ and spectral matching, where pre-recorded MS/MS spectra of known compounds from pure standards are compared to experimental spectra to identify matching fragments^7^. The success of this method is heavily dependent on the availability of comprehensive and accurate MS/MS spectral libraries. Unfortunately, current spectral libraries suffer from limitations in size, quality, and diversity, and they are difficult to maintain and update^8^,^9^. As a result, a substantial portion of MS/MS spectra generated in metabolomic experiments remains unidentified through spectral library searching methods.

To address the challenge of metabolite identification, computational tools generally adopt one of two strategies^10^: (i) computationally processing experimental MS/MS spectra and searching for putative annotations among molecules in databases of known compounds^11,12^; or (ii) using the chemical structures of metabolites to predict their MS/MS spectra^13–18^, which are then compared to experimental spectra for identification. Both approaches involve machine learning algorithms that convert experimental MS/MS spectra into feature vectors and encode chemical structures as fingerprints, embeddings, or graph structures for learning purposes.

In this study, we focus on the second approach, which aims to overcome the limitations of empirical MS/MS libraries by computationally predicting MS/MS spectra, ultimately generating reliable *in silico* MS/MS libraries. Existing tools that follow this approach employ a combination of techniques: rule-based methods that capture known fragmentation patterns based on chemical principles^13,16^, probabilistic models that assign probabilities to potential fragmentation events through statistical analysis of experimental MS/MS spectra^14,15^, and machine learning algorithms, ranging from traditional methods to deep learning^17–19^.

Here, we introduce SingleFrag, a novel deep learning tool for predicting MS/MS spectra (Fig. 1). SingleFrag stands out from existing machine learning approaches in two significant ways. First, instead of predicting the entire fragmentation spectrum of a molecule with a single model, we train a separate model for each fragment ion. This approach acknowledges that peaks (corresponding to fragment ions) in an MS/MS spectrum are not necessarily correlated. For example, one peak might correspond to a particular moiety in the molecule, while another peak corresponds to a different moiety with no relation to the first. Using the same features to predict both peaks simultaneously can reduce prediction accuracy. Therefore, by training individual models for each fragment, we aim to better capture the molecular and structural features associated with each specific peak. Second, rather than predicting the intensity of peaks in the spectrum, SingleFrag focuses on predicting the presence or absence of each peak. We argue that the presence of a given fragment is more relevant for molecular structural annotation than its intensity, which can vary based on factors such as the fragmentation technique (for example, collision-induced dissociation or high-energy C-trap dissociation), collision energy, and even the mass spectrometer manufacturer. This approach allows for more robust and reliable predictions across different experimental conditions.

**Figure 1.**
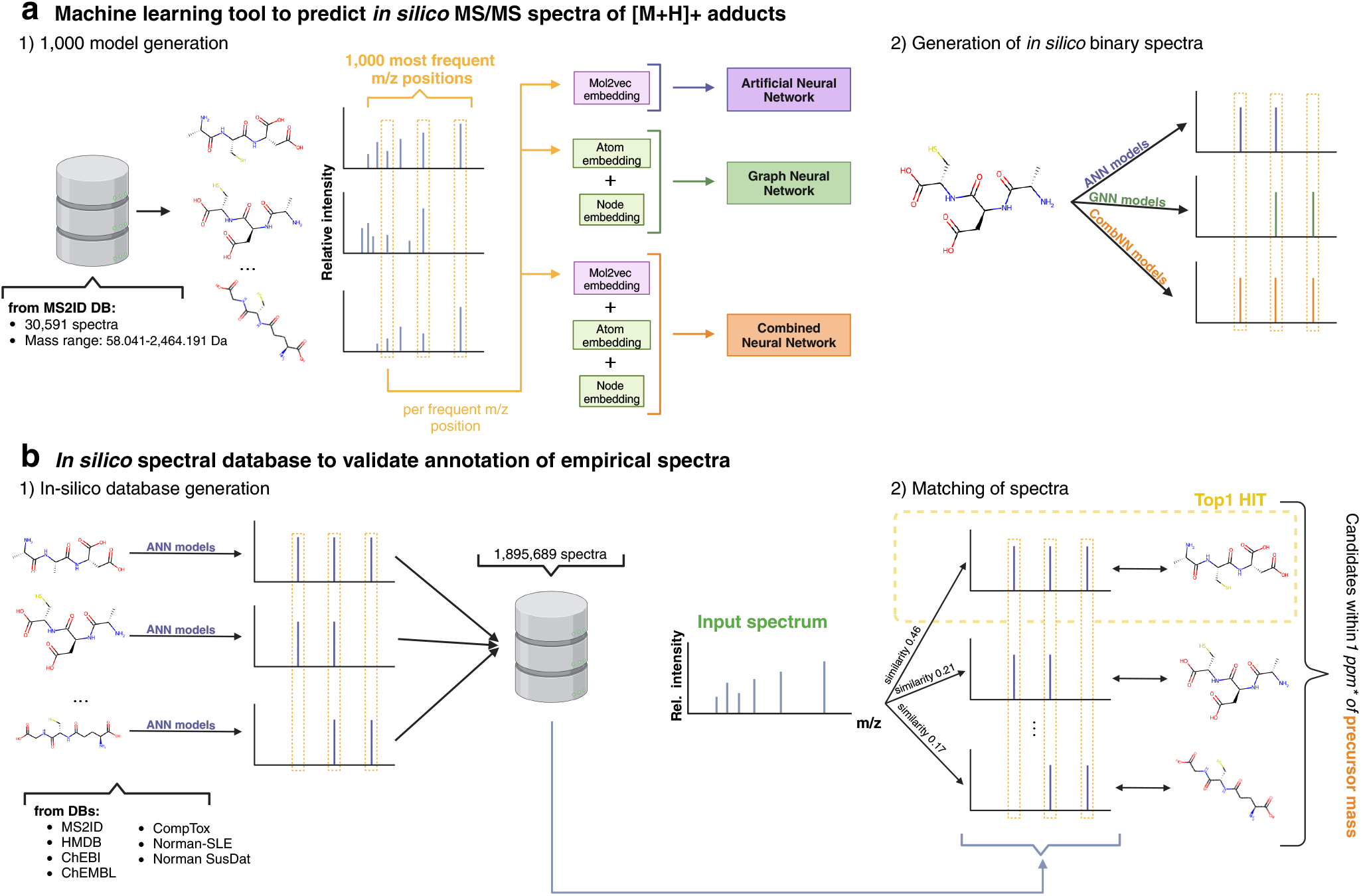
SingleFrag for spectral prediction and annotation. **a**, We consider an empirical database of MS/MS spectra, and train three model types to predict the presence or absence of each of individual fragment ions across spectra. In particular, we build 1,000 models (of each of the three model types), corresponding to the 1,000 most frequent fragment ions. Once all models are trained, we predict the whole spectrum of a given molecule by predicting each of the 1,000 peaks independently. **b**, To annotate unknown empirical spectra, we build a database of *in silico* predicted spectra for over 1.8 million compounds. Then, for an unknown empirical spectrum that we wish to annotate, we select all candidates from the database with masses compatible with the unknown spectrum, and rank the candidates according to the similarity between their predicted spectra and the target empirical spectrum.

We train three SingleFrag models to predict the presence of individual fragments (Fig. 1a). The first model embeds the molecule using mol2vec^20^ and uses this embedding as input to a multilayer, feedforward neural network. The second model employs a graph neural network (GNN)^21–23^ that takes as input the unprocessed molecular graph. The third model integrates both the mol2vec embedding and the graph neural network. We find that all three models, particularly the first one, outperform state-of-the-art methods at predicting MS/MS fragments, including rule-based (CFM-ID^16^) and deep learning models incorporating the 3D structure of the molecule (3DMolMS^19^). Encouraged by this performance, we generate an *in silico* MS/MS library for nearly 2 million molecules and test its ability to identify unknown metabolites from their spectra (Fig. 1b). For each target spectrum, we rank candidates from the library based on the similarity between the experimental spectrum and SingleFrag’s *in silico* predictions. We find that the true target molecule is ranked first in 38% of cases and within the top five candidates in 72% of cases, demonstrating high accuracy and reliability. Finally, we apply this annotation method to recurrent unidentified spectra from the ARUS database^24^ of the NIST Mass Spectrometry Data Center, successfully annotating three metabolites. We confirm these annotations by manually analyzing their molecular structures and fragmentation patterns, thereby validating the effectiveness of our approach.

## Results

### SingleFrag models for *in silico* prediction of individual MS/MS fragments

We develop and validate our models using a dataset containing MS/MS fragmentation spectra for 30,591 compounds sourced from HMDB^25^, Agilent METLIN, MassBankEU and MoNA^26,27^, NIST, Riken^28^ (Methods). We allocate spectra from 24,473 of these compounds (80% of the total) to the training set, while the remaining compounds are evenly divided into validation and test sets. All results reported here correspond to the test set, which is not used in any way for training or hyperparameter tuning. In all cases, we discretize the spectra into bins of size m/z = 0.01 by ceiling the mass of each fragment to two decimal places.

Instead of building a single model to predict the entire MS/MS spectrum of compounds, we construct a separate model for each mass bin. This approach typically results in models for individual fragments, hence the name SingleFrag (however, some bins contain distinct fragments; see Methods and Supplementary Figure 1). Building a model for every bin computationally very demanding, so we focus on the 1,000 bins where peaks occur most frequently, thereby concentrating resources on the most informative and data-rich parts of the MS/MS spectra. These bins cover masses m/z in the range [29.04, 269.09], and account for 60% of all peaks in the spectra in our dataset (Fig. 2a). By restricting spectra to these 1,000 bins, 94% of the compounds in the dataset still have 3 or more peaks, 90% have 5 or more, and 80% have 10 or more (Fig. 2b).

**Figure 2.**
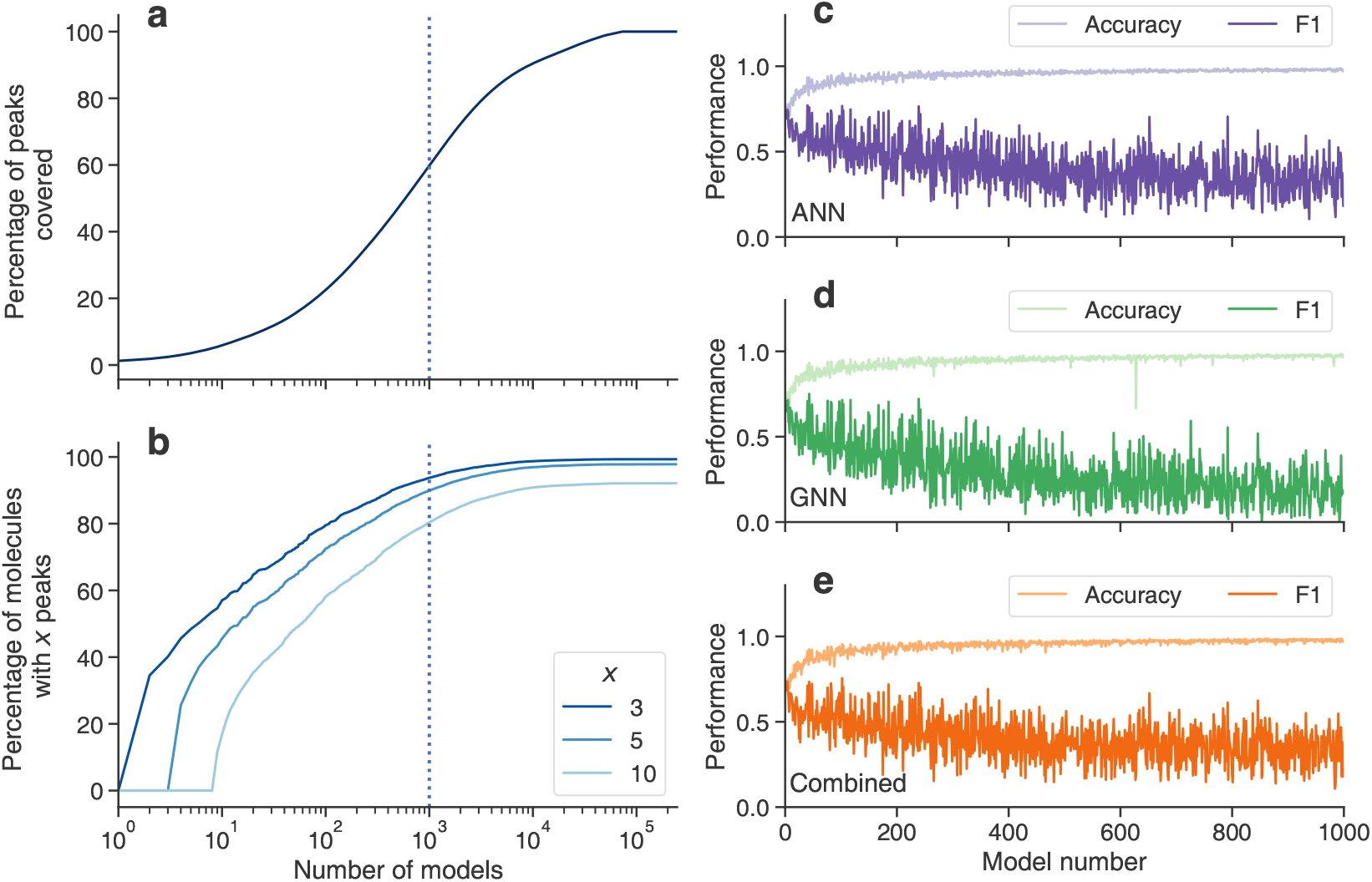
Prediction of individual fragments. We discretize the m/z axis of each spectrum into bins of width 0.01. (**a**) The percentage of peaks in the training set covered by modeling only the x bins with the highest frequency of peaks across molecules (x-axis). The dashed line indicates the coverage achieved by 1,000 models. (**b**) The percentage of molecules in the training set whose spectra contain 3, 5 and 10 peaks when only the *x* most frequent bins are considered. The dashed line indicates the coverage achieved by 1,000 models. Based on (**a**) and (**b**), we select the 1,000 bins that cover the highest fraction of peaks and molecules. These bins (**a**) cover 60% of the peaks in the training set and (**b**) ensure that 94% of the training spectra have at least 3 peaks, 90% have at least 5 peaks, and 80% have at least 10 peaks. For each of these bins, we train three different SingleFrag models (ANN, GNN, and combined; see text for details). (**c** to **e**) Accuracy and *F*_1_ score for the each bin and model type: (**c**) ANN, (**d**) GNN, and (**e**) combined. The models on the *x*-axis are sorted from highest to lowest frequency of peaks in the training set, with the first model corresponding to the bin that most frequently contains a peak (at m/z 51.03) in the training set.

Each of the 1,000 models is a binary classifier that takes as input the molecule and predicts whether a peak exists in the corresponding mass bin. We focus on binary predictions because the presence of a given fragment is more relevant for molecular structure annotation than its intensity, which can vary based on factors such as collision energy or the mass spectrometer used to obtain the spectrum. To account for all possible fragments associated with a molecule, we consider all available MS/MS spectra for a given molecule and build a single merged discretized spectrum with binary values, so that m/z bins in which a peak is present in at least one spectrum are equal to one, and other bins are equal to zero (Methods).

We investigate three different SingleFrag models as our peak binary classifiers (see Methods for details). First, we use a multilayer feedforward artificial neural network (ANN). In this model, molecules are embedded into a 300-dimensional space using the previously trained mol2vec model^20^, and the resulting embeddings are fed into the neural network. Second, we employ a graph neural network (GNN) that directly uses the molecular graph as input^21–23^. Finally, we combine these approaches into a model that uses both the graph neural network and the mol2vec embeddings. Each SingleFrag model returns a score between 0 and 1 for each input molecule and m/z bin. To evaluate the model’s performance, we convert these scores into binary predictions for the presence (1) or absence of a peak. To this end, we calculate a specific threshold for each m/z bin, so that scores above the threshold are converted into a 1 and scores below the threshold are converted into to a 0. We select this threshold so that predictions are well-calibrated, that is, that the fraction of molecules predicted to have a specific peak matches the fraction of molecules with that peak in the training set (see Methods for details).

Fig. 2c-e illustrates the performance of each model type (ANN, GNN, and combined) across different mass bins. We evaluate performance using accuracy and the F1 score (see Methods for details). Due to the unbalanced nature of the target values—where even the most frequent peaks are relatively rare—accuracy often approaches 1 and does not provide a clear picture of model performance. In contrast, the F1 score, which is close to both precision and recall because our models are well-calibrated, is more informative. Our results show that the ANN and combined models, with mean F1 scores of 0.411 and 0.399 respectively, outperform the GNN model, which has a mean F1 score of 0.290.

### SingleFrag models yield accurate *in silico* spectra

Next, we investigate whether SingleFrag models, trained to predict individual peaks, can accurately predict whole spectra. For this, we apply all 1,000 bin-specific models to each molecule in the test set to obtain the corresponding predicted spectrum (Fig. 3). We perform this analysis for each type of SingleFrag model (ANN, GNN, and combined; Fig. 3c-e). To evaluate the performance of the SingleFrag models, we benchmark them against two state-of-the-art algorithms: (i) CFM-ID^16^ (Fig. 3b), the leading rule-based prediction algorithm, and widely used for *in silico* fragmentation; and (ii) 3DMolMS^19^ (Fig. 3a), which takes into account the 3D structure of molecules in its predictions. We compare the predicted spectra in terms of precision, recall, accuracy, and cosine similarity between the predicted and real spectra in the test set (Fig. 3g-i). These performance metrics vary considerably across compounds. Given that our validation naturally provides matched samples (predictions from different algorithms for each compound), we report the number of molecules for which each tool has the best metric. Additionally, we restrict the comparison to molecules that could be predicted by all methods, specifically those for which both CFM-ID and 3DMolMS return valid outputs, accounting for 2, 929 out of 3, 059 compounds in the test set. To ensure consistency, we set the size of m/z bins to 0.2, matching the precision provided by 3DMolMS. As before, we disregard the intensity of fragment ions and focus solely on whether peaks are predicted to exist or not.

**Figure 3.**
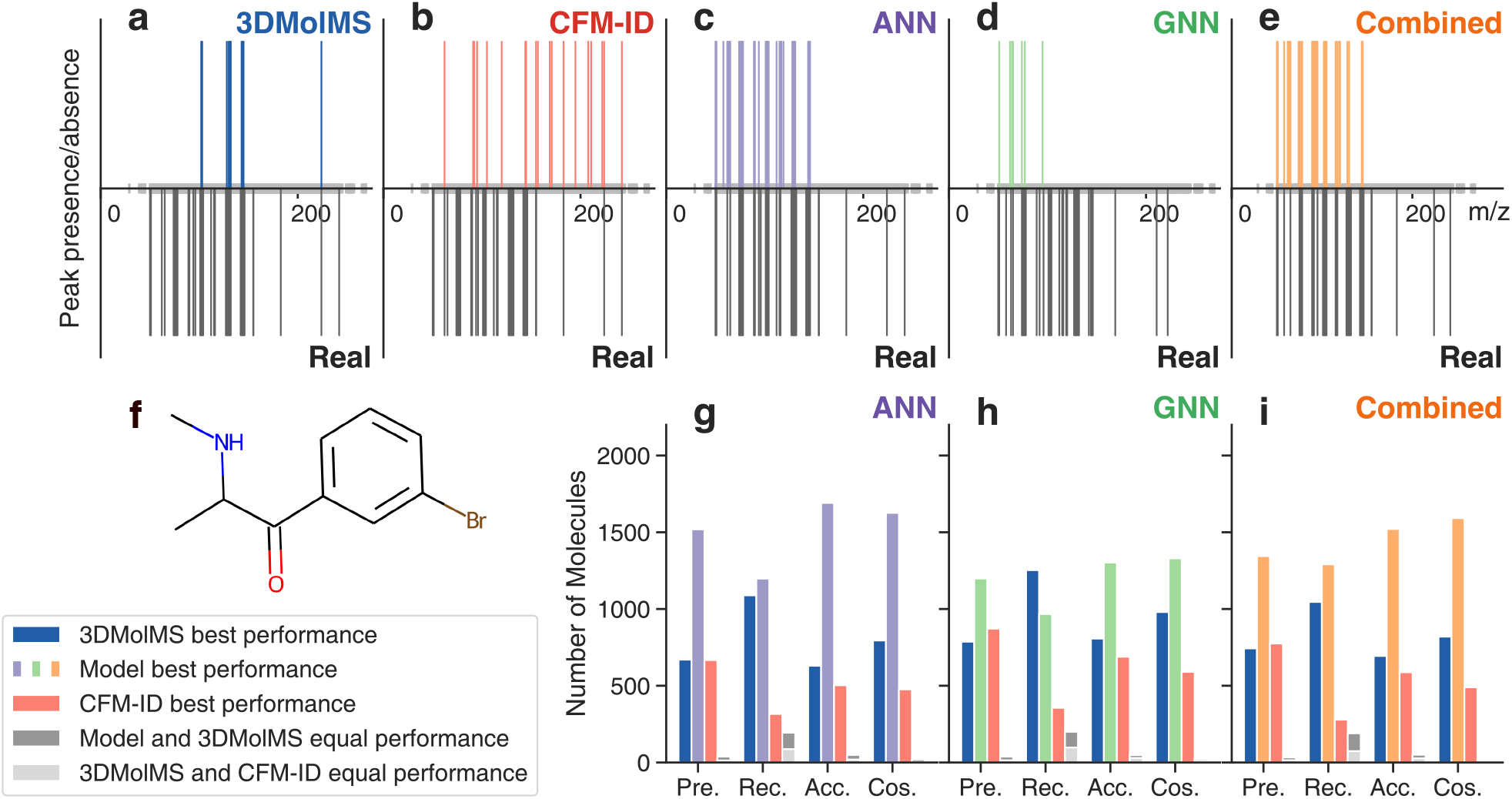
Prediction of whole spectra. (**a**-**e**) We predict the MS/MS spectrum of the molecule depicted in (**f)** using: (**a**) 3DMolMS^19^; (**b**) CFM-ID^16^; (**c**) SingleFrag ANN model; (**d**) SingleFrag GNN model; and (**e**) SingleFrag combined model. As throughout the paper, we disregard the intensity of peaks and just represent their presence or absence. Note that, by construction, SingleFrag models cannot predict all peaks, but only those at the 1,000 most frequent bins (grey ticks in the m/z axis; see text for details). However, when comparing to the real spectra, all m/z positions are considered. (**g**-**i**) We benchmark the performance of SingleFrag models against 3DMolMS and CFM-ID. For each molecule in the test set, we calculate the precision, recall, accuracy, and cosine similarity of the predicted spectrum compared to the real spectrum. Since these performance metrics vary considerably across molecules, and given that our validation naturally gives matched samples (that is, for each molecule we obtain the spectrum predicted by each of the algorithms), we plot, for each metric, the number of molecules for which a given model is the best performer.

As shown in Fig. 3g-i, we find that 3DMolMS performs better than CFM-ID, as expected^19^. Although both algorithms perform similarly in terms of precision, 3DMolMS tends to have higher recall, resulting in higher cosine similarities between the predicted and real spectra. However, all three SingleFrag models achieve better precision than both CFM-ID and 3DMolMS. Since their recall is comparable to 3DMolMS (slightly better for the ANN and combined models, and slightly worse for the GNN model), SingleFrag models produce *in silico* spectra with higher accuracy and cosine similarity to the true spectrum overall. This is particularly noteworthy considering that SingleFrag cannot predict peaks outside the 1,000 trained m/z bins. The fact that, despite this limitation, the overall predicted spectrum is closer to the true spectrum confirms our hypothesis that modeling individual peaks separately results in more accurate predictions than using a single model for the entire spectrum.

### A database of *in silico* spectra of known molecules enables the annotation of unknown empirical spectra

Given the success of SingleFrag models, particularly the ANN model, in predicting whole spectra, we investigate whether these *in silico* predicted spectra are accurate enough to annotate unknown MS/MS empirical spectra of known compounds. These are compounds that have been described in chemical databases but whose fragmentation spectra are not available to the metabolomics community. To facilitate this annotation process, we have created a database containing nearly 1.9 million *in silico* MS/MS spectra predicted by the ANN SingleFrag model. To generate these spectra, we begin with the SMILES representations of almost 1.9 million compounds (Methods). Using these SMILES, we first obtain their 300-dimensional mol2vec embeddings, and then perform a forward pass through the SingleFrag ANN model to predict their spectra.

With the resulting database, we can annotate unknown spectra through the following steps. Given an unannotated empirical spectrum, we first estimate either: (i) the molecular mass of the corresponding compound based on the mass of the precursor ion, or (ii) its exact molecular formula using a tool such as BUDDY^29^. Next, we filter the database of 1.9 million compounds/spectra to find candidate compounds that match the mass or molecular formula, respectively, of the unknown target compound. Finally, we rank these candidates by calculating the cosine similarity between the unannotated empirical spectrum and the *in silico* spectrum predicted for each candidate.

To validate the performance of SingleFrag, we use the same test set of molecules as in previous sections. For each MS/MS spectrum, we generate a list of candidate molecules with compatible molecular formulas (Fig. 4a,d) or compatible molecular masses, assuming a spectrometer precision of either 1ppm (Fig. 4b,e) or 10ppm (Fig. 4c,f). We evaluate the annotation performance by tracking the position of the true molecule in the candidate ranking for each test molecule. In Fig. 4a-c, we show the frequency with which the true molecule is ranked first (Top 1, indicating perfect annotation), among the top five candidates (Top 5), and among the top ten candidates (Top 10). We measure this performance under four experimental conditions: using low (0-14 eV), medium (15-39 eV), or high (*≥* 40 eV) collision energy spectra as the unknown empirical query (800, 1,057, and 542 spectra, respectively, from the NIST20 and Agilent METLIN Metabolomics databases; Methods), and using merged query spectra where peaks at all available collision energies are combined into a single spectrum (see Methods for details).

**Figure 4.**
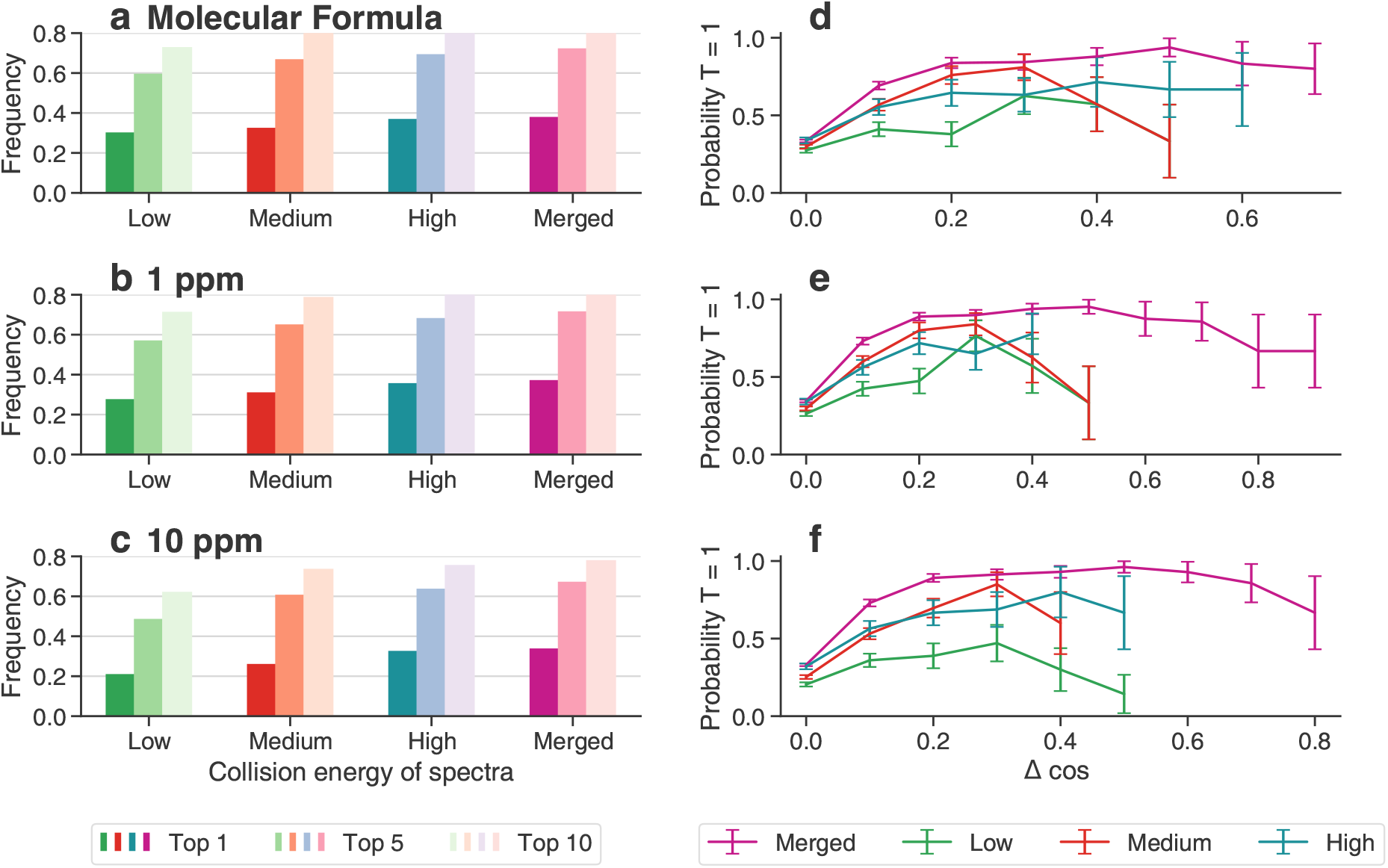
Annotation of unknown empirical MS/MS spectra. We evaluate the ability of SingleFrag to annotate empirical MS/MS spectra of known compounds, meaning compounds that are described in databases but whose reference fragmentation spectra are not available. For each molecule in the test set, we identify putative annotations by selecting compatible candidates from a reference database containing nearly 1.9 million known molecules. Compatibility is established based on: (**a, d**) exact molecular formula (simulating a scenario where the exact formula of the test molecule is determined with a tool such as BUDDY^29^); (**b, e**) exact mass, with a window of ±1 ppm; or (**c, f**) exact mass, with a window of ±10 ppm. We then rank candidate annotations by computing the cosine similarity between the empirical spectrum of the test molecule and the predicted SingleFrag ANN spectrum of each compatible candidate. For each test molecule and experimental condition (exact formula, *±* 1 ppm, and *±* 10 ppm), we consider separately empirical spectra with different collision energies: (i) low collision energy (*<*15 eV); (ii) medium collision energy (*≥*15 eV and *<*40 eV); (iii) high collision energy (*≥* 40 eV). Additionally, we consider merged spectra processed as in the training of SingleFrag, by merging spectra for all collision energies available for each test molecule. (**a**-**c**) We plot the frequency with which the true molecule (Tanimoto coefficient^30^ *T* = 1) is ranked as the top candidate (perfect annotation), and among the top 5 and top 10 candidates (plausible annotations). (**d**-**f**) For each experimental condition, we plot the probability of the top candidate being the true molecule (Tanimoto coefficient^30^ *T* = 1) as a function of the score gap Δcos between the top and the second ranked candidates. Generally, larger gaps provide more confidence in the top annotation.

We obtain the best results when we use as much information as possible, that is, when the exact molecular formula of the target molecule is assumed to be known from a tool such as BUDDY^29^, and multiple spectra (at different collision energies) for the molecule are merged (Fig. 4a). Under these conditions, the true molecule is ranked first in 37.6% of cases, among the top five candidates in 71.9% of cases, and among the top ten candidates in 82.7% of cases. Performance decreases only slightly when individual high-energy spectra are used (36.6% ranked first, 69.0% top five, 81.2% top ten). Performance decreases more noticeably when medium-energy spectra (32.2% ranked first, 66.5% top five, 79.9% top ten) and low-energy spectra are used (29.9% ranked first, 59.3% top five, 72.6% top ten). This happens because SingleFrag focuses on the 1,000 bins where peaks occur most frequently, covering masses in the low mass range between m/z 29.04 and 269.09. These bins are more likely to be present in high-energy spectra, which typically produce extensive fragmentation, especially in the low mass range. By contrast, low-energy spectra tend to be less fragmented, dominated by a few large fragments or the molecular ion itself, for which SingleFrag makes no predictions if the molecules have larger m/z than 269.09.

Significantly, in all cases, the reliability of annotations can be considerably increased by examining the relative position of the candidates in the ranked list (Fig. 4d-f). As mentioned, we score candidates based on the cosine similarity between the empirical spectrum of the target metabolite and the *in silico* predicted spectrum for each candidate. However, the difference between the scores of the first and second candidates is particularly informative about the reliability of the top candidate. When the two scores are very close, the top candidate is the true metabolite in 20-30% of cases (Fig. 4d-f), consistent with the overall figures in Fig. 4a-c. However, when there is a large gap between the cosine similarities of the first and second candidates, the top candidate is much more likely to be the true molecule. Specifically, when the difference in cosine similarity scores (Δcos) between the top and second ranked candidates exceeds 0.2, the top candidate is the correct molecule in the majority of cases. (This is specially true for merged spectra and at high collision energies; at low energies, the small number of instances with gaps larger than 0.4 sometimes obscures this signal.)

### Annotation of recurrent unidentified spectra from the ARUS database

Building on the validation results from the previous section, we demonstrate that our approach effectively annotates unknown spectra. We focus on spectra from the Annotated Recurrent Unidentified Spectra (ARUS) database^24^, maintained by the NIST Mass Spectrometry Data Center. This database includes spectra that frequently appear in real samples but remain unannotated.

We used a dataset of ARUS spectra with putative molecular formulas assigned using BUDDY^29^. The dataset includes spectra from two sources: plasma (25,801 spectra) and urine (68,478 spectra). For each unknown molecule, we discretized and binarized its associated spectrum as previously described. We then generated a list of candidate annotations for each spectrum using a filtering window of 1 ppm. To be as exhaustive as possible in the annotation of these unidentified spectra, we enlarged the *in silico* database with all compounds available for download from PubChem, resulting in an extended database of 96,492,904 predicted spectra. We ranked the candidate annotations using the similarity between the target empirical spectrum and the SingleFrag *in silico* prediction for each candidate’s spectrum and kept the ten candidates whose *in silico* spectra were more similar to the query spectrum.

From hundreds of potential annotations, we selected three particularly promising ones. These selections were based on two criteria: (i) the molecular formula matched BUDDY’s prediction, with a low estimated false discovery rate in BUDDY; and (ii) there was a significant gap between the first and second candidates in SingleFrag’s ranking. We confirmed the annotation for these three compounds—glucuronyl-2-hydroxyhippurate, 8-hydroxyquinoline glucuronide, and a truxilline isomer—through the manual elucidation of their fragment ions from their molecular structures by an expert chemist (Fig. 5). These three compounds are of particular interest due to their putative origin. Isomeric truxillines, a group of minor alkaloids consistently found in illicit cocaine samples, suggests possible exposure through drug use or environmental contamination related to cocaine production. 8-hydroxyquinoline glucuronide, a metabolite of 8-hydroxyquinoline, could indicate pharmaceutical use or exposure to quinoline compounds. Glucuronyl-2-hydroxyhippurate, a conjugate of hippuric acid, may reflect dietary intake or endogenous metabolic processes involving aromatic acids.

**Figure 5.**
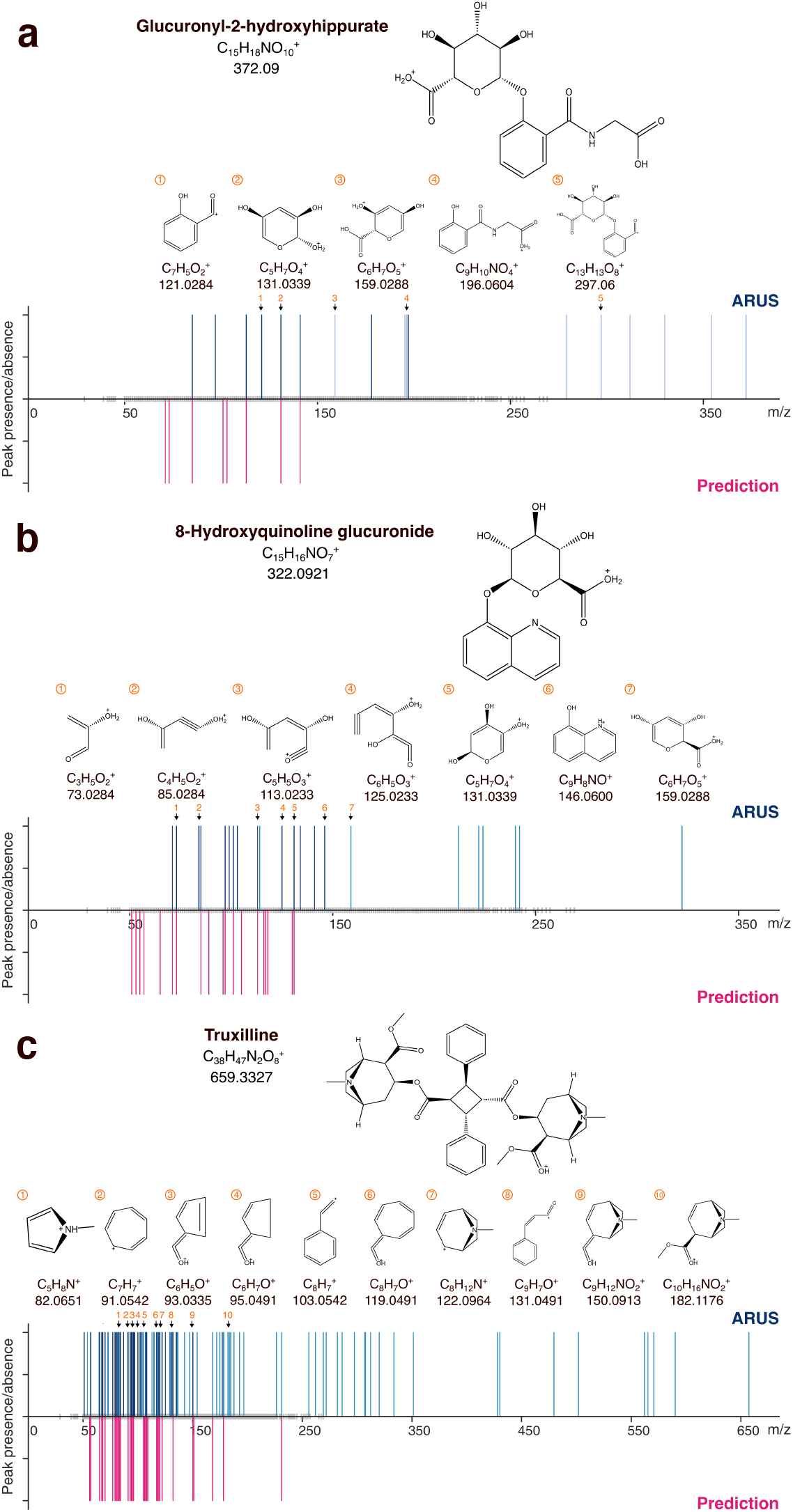
Annotation of three empirical MS/MS spectra from the ARUS database24. For three different unannotated spectra in ARUS, we show the empirical MS/MS spectrum (obtained by merging all available spectra for the compound and binarized as in the rest of the paper) and the predicted spectrum for the top candidate annotation identified by SingleFrag. In each case, we show the chemical structure of the top candidate, as well as the match between observed peaks of the empirical spectrum and plausible fragments identified manually via structural analysis of the molecule.

## Discussion

SingleFrag is a novel *in silico* fragmentation tool for metabolites and small molecules. Unlike existing machine learning tools, SingleFrag focuses on predicting the presence or absence of individual fragments rather than attempting to predict whole spectra, including all peaks and their intensities. This approach is based on the rationale that the molecular features predictive of the existence of a specific fragment may be distinct from those predictive of other fragments. Our results validate this rationale.

We developed three different SingleFrag models: ANN, GNN, and combined. These models operate in fundamentally different ways. The ANN model uses a mol2vec embedding, while the GNN model works directly with the molecular graph, leading to distinct neural network architectures and underlying mathematical models. Despite these differences, all SingleFrag models outperform the state-of-the-art *in silico* prediction tools, CFM-ID and 3DMolMS. This is particularly noteworthy for 3DMolMS, which employs a more sophisticated network architecture and utilizes detailed features encoding the 3D location of each atom in molecules.

We hope that our approach of modeling peaks individually will inspire new developments where more complex features and models, such as those used in 3DMolMS, can be integrated into SingleFrag-like frameworks to achieve even higher levels of spectral predictive accuracy. Additionally, we recognize that there is significant potential for enhancing the neural network architectures we have employed. Indeed, we deliberately kept our models simple to demonstrate that the improved performance is primarily due to the approach of modeling individual peaks instead of whole spectra, rather than more sophisticated deep learning techniques. This is especially relevant for the GNN model. Its slightly lower performance compared to the ANN model should not be seen as a fundamental limitation of graph neural networks in the domain of spectral prediction, but rather as an opportunity to refine graph features and optimize graph layer architectures for even better results.

We have also shown that, even with relatively simple neural network models, SingleFrag can already be used to annotate real spectra, both in a controlled setting with test molecules, and in a real-world scenario with ARUS molecules that we have annotated for the first time. We expect that future SingleFrag-like algorithms will enable even more reliable annotation, which is the standing roadblock for the development of the full potential of metabolomics.

Moreover, our results emphasize the critical importance of acquiring experimental MS/MS spectra for each precursor ion at multiple collision energies. Among the widely used spectral libraries, only the NIST MS/MS provides a systematic and curated repository of spectra acquired at multiple collision energies. By contrast, other spectral libraries such as GNPS, MSDial, or MassBank often lack sufficient curation and typically contain spectra acquired at only one or a few collision energies per compound. This limitation can lead to incomplete or less informative spectra, which may not fully represent the diversity of fragment ions produced under different collision conditions.

By employing techniques like stepped collision energy, it is possible to capture a broader range of fragment ions, resulting in more detailed and informative spectra. Our results suggest that this approach may be essential, not only for real metabolomic experiments, but also for the expansion and curation of existing reference spectral libraries. This creates a positive feedback loop—improved predictive models generate better *in silico* spectra, which in turn can more accurately match the experimental spectra of unidentified compounds. This enhances the reliability and utility of these predictions, ultimately leading to new discoveries and a more comprehensive understanding of metabolic pathways and processes.

## Methods

### Machine learning tool to predict *in silico* MS/MS spectra of [M+H]+ adducts

**Data** We obtained 2,291,119 MS/MS spectra from the HMDB, Agilent METLIN, MassBankEU and MoNA, NIST, and Riken databases, corresponding to 478,631 unique compounds. The annotations for these spectra and compounds may include the name, molecular formula, InChIKey, SMILES, adduct, precursor ion mass, collision energy, and mass spectrometer type.

After excluding *in silico* generated spectra (mainly from HMDB), we retained 450,248 experimental MS/MS spectra for the protonated adduct in positive mode ([M+H]). We discarded spectra with m/z values reported with a single decimal or generated by low-resolution instruments (see Supplementary Material for details). After filtering, we had 434,480 spectra corresponding to 54,790 compounds. To ensure consistency, we obtained canonical SMILES from the latter via their InChIKeys and the Chemical Identifier Resolver API service of PubChem (https://pubchem.ncbi.nlm.nih.gov/rest/pug/compound/inchikey/). After removing 11,916 molecules without InChIKeys and 12,282 molecules not found by the request, we obtained a final dataset of 30,591 unique compounds.

We then merged all spectra corresponding to each unique compound into a single spectrum, regardless of the original database or acquisition tool. Using unique database identifiers for each molecule, we combined the m/z values from all corresponding spectra into a single binary list (see below details about how we build binned spectra).

We ensured that each molecule in the dataset had its exact mass, verifying that it matched the mass calculated from its SMILES within a 0.01 difference. Only one molecule did not match, so we eliminated it to avoid inconsistencies. We also verified that each spectrum contained the m/z of the precursor ion corresponding to the [M+H] adduct, allowing a window of *±* 0.05 Da. For spectra without such a precursor ion, we manually added it by summing the mass of a proton to the neutral mass of the compound. Additionally, we removed the peaks for m/z values larger than the mass of the precursor ion plus 0.05. This process resulted in a dataset with molecules having masses between 58.041 and 2,464.191 Da. Finally, we randomized the data and divided it into a training set containing 80% of the molecules (24,472 spectra), and validation and test sets containing 10% each (3,059 spectra each).

### Spectrum binarization and merged spectrum construction

SingleFrag predicts the existence/absence of peaks in m/z bins with a resolution of 0.01 m/z. To that end we discretize the 450,248 spectra in our database by assigning peaks to the m/z bin corresponding to the ceiling of the second decimal of the m/z value corresponding to the peak. For instance, if a spectrum has a peak (fragment) at m/z 110.253, we assign it to m/z bin 110.26. Because, we binarize spectra, m/z bin 119.26 would have associated a value of 1 since there is a peak in that bin. At the end of the process, the binarized, discrete spectrum is a vector in which a value of 1 indicates the presence of a peak in the corresponding bin and a value of 0 indicates the absence of a peak in that m/z bin.

To obtain the merged spectrum of each unique compound, we consider all individual spectra for that compound. For each m/z bin, we look at whether individual spectra have a peak in that bin. If at least one spectrum has a peak, we assign a 1 to that bin, and a 0 otherwise. Therefore the merged spectrum of a compound represents all the possible fragments that have been detected for that compound.

Note that binning spectra is a necessary choice when using machine learning methods with discrete inputs. As explained in the main text, by using 0.01 m/z bins and using the 1,000 most commons bins with associated fragments, we have a good coverage of the fragments observed in the majority of spectra. We note that, by binning spectra into 0.01 m/z bins, nonzero intensities associated to the same fragment may be split into two neighboring bins—for instance, fragments with m/z values 25.889, 25.891 would be classified in different bins, but could have been attributed to the same bin had we binned spectra differently. To assess how often this happens, we chose 100 random bins of size m/z 0.01 and represented the distribution of fragments associated to that bin (Supplementary Figure 1). Our results show that approximately half of the time the fragments were distributed in the middle of the bins, and in the other half the fragments were distributed between neighboring bins, which suggests that there is no perfect binning strategy.

### Machine learning models

We introduce three SingleFrag models. All models aim to predict whether a molecule has a peak at a specific mass bin or not, and are thus binary classifiers:

#### 1. Artificial Neural Network (ANN)

First, we create 300-dimensional embeddings for all molecules in the training, validation and test sets using the mol2vec algorithm^20^. These embeddings were then input into a Multilayer Perceptron with three fully-connected layers, two of them with ReLU activation functions, and the final layer with a Sigmoid activation. The model was trained with a batch size of 16, a learning rate of 1*×* 10^*−*4^, and binary cross entropy as the loss function. We set a minimum of 200 and a maximum of 2,000 epochs for training, and keep the model with the lowest validation loss. We also implemented an early auto-stop if the validation loss increased while the training loss decreased, comparing the most recent epoc to the previous 100 epochs.

#### 2. Graph Neural Network (GNN)

The GNN model uses the whole molecular graph to make predictions. We used bond embeddings (type of bond and atom indices) and atom embeddings (neighboring atom sequence, total hydrogens, formal charge, mass, and aromaticity). We run the node embeddings through three graph attention layers (GAT; these assign different importance to each connection in the graph) with ReLU activations. After the third GAT layer, we aggregated embeddings using sum, max, and average functions, and used a sigmoid as an activation function. We used training parameters equal to those of the ANN model but with a minimum of 4,000 and a maximum of 10,000 epochs.

#### 3. Combined Neural Network (CNN)

This model integrates the ANN and GNN models by using both the 300-dimensional mol2vec embeddings and the graph embeddings. The first three layers of the CNN matched the GNN structure, using graph attention operators and ReLU activations, followed by pooling. To integrate the GNN and ANN predictions, we reshaped the mol2vec vector dimensions to align with the pooled data. The ANN component included a multilayer perceptron with three fully-connected layers, two ReLU activations, and one sigmoid activation. After combining these outputs, we applied a linear layer with 45 neurons and another linear layer with ReLU, followed by a final linear layer with a sigmoid activation. Training parameters were consistent with the previous models, with a batch size of 16, a learning rate of 1e-4, binary cross entropy loss, and a training range of 4,000 to 10,000 epochs.

### Threshold scores and model calibration

To produce *in silico* binary spectra, we need to convert the output scores from our machine learning models (ANN, GNN and CNN) into a 0 or 1 prediction for the existing of each peak. A way to achieve this is to specify a threshold value for each model associated to a m/z bin so that scores above the threshold are converted into a 1 (presence of a peak) and the remaining scores are converted into a 0 (absence of a peak).

Because the training and test sets are selected at random, the expected fraction of molecules having a peak in a specific m/z bin in the train, validation and test sets is the same. Therefore we have to select a threshold that recovers the statistically correct fraction of molecules with peaks in that bin. To that end, for a specific m/z bin, we rank the *N*_val_ molecules in the validation set according to their score for that bin. We then compute the fraction *f*_m*/*z_ of molecules with a peak in that bin within the training set. We set the threshold as the score of the molecule in position *f*_m*/*z_*N*_val_ within the validation set (rounded upwards to the next integer). Note that the threshold value depends on the modeling approach (ANN, GNN and CNN) and the m/z bin we consider. By binarizing scores in this way, we ensure that our models are statistically calibrated.

### Validation

To evaluate our predictions, we compare them with those of two other *in silico* fragmentation tools: CFM-ID^16^ and 3DMolMS^19^.

CFM-ID uses competitive fragmentation modeling and machine learning to fit model parameters. The latest version, CFM-ID 4.0, is accessible via a web server at https://hub.docker.com/r/wishartlab/cfmid ^16^. Using CFM-ID 4.0 with its default values, which exclude fragmentations below a 0.001 probability threshold, we obtained predictions for 3,034 out of the 3,059 molecules in the test set. The tool CFM-ID 4.0 calculates spectra for low (10 eV), medium (20 eV), and high (40 eV) collision energies, representing them as lists of mass-intensity pairs, each corresponding to a peak in the spectrum. We then consolidated the spectra predicted at different collision energies into a single merged spectrum.

3DMolMS uses deep learning to predict MS/MS spectra from 3D molecular conformations and other molecular features. The source code for the tool is available from https://github.com/JosieHong/3DMolMS. We used the pretrained model provided in this repository, and generated *in silico* spectra at low (10 eV), medium (20 eV), and high (40 eV) collision energies. We then consolidated the spectra predicted at different collision energies into a single merged spectrum. 3DMolMS produces spectra at 0.2 m/z resolution.

Ultimately, 2,929 spectra were predicted using CFM-ID 4.0, 3DMolMS, and SingleFrag. Because 3DMolMs produces spectra at a 0.2 m/z resolution, we discretized CFM-ID 4.0 and SingleFrag spectra at the same resolution. We also consider only binary spectra and do not use information about the intensities associated to m/z bins or fragments.

To evaluate our models, we calculated various metrics comparing the real MS/MS spectra of the molecules (discretized and binarized) with their *in silico* spectra reconstructed by different methods (ANN, GNN, Combined, CFM-ID and 3DMolMS). The metrics used were precision, recall, accuracy, F1 score, and cosine similarity. To define these metrics, we use the following nomenclature: true negative (TN), true positive (TP), false positive (FP), and false negative (FN). A model prediction is Positive if it is 1 and Negative if it is 0. Additionally, a prediction is True if it is correct and False if it is incorrect.

- **Precision**: Measures the proportion of true positive predictions among all positive predictions made by the model. It indicates how many of the predicted positives are actually correct.

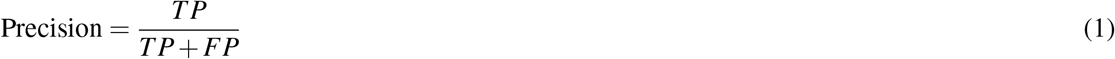
- **Recall**: Also known as sensitivity, it measures the proportion of true positive predictions among all actual positives. It indicates how well the model can identify positive instances.

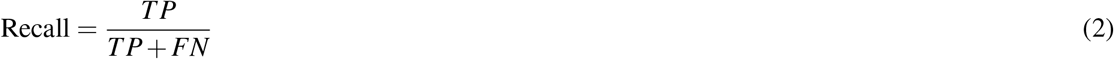
- **F1**: Combines Precision and Recall into a single metric by taking their harmonic mean. It provides a balance between Precision and Recall, especially useful when there is an uneven class distribution.

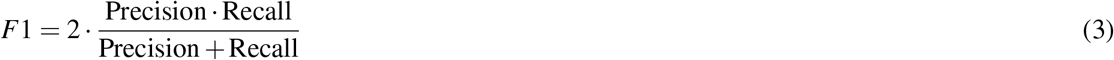
- **Accuracy**: Measures the proportion of all correct predictions (both true positives and true negatives) among the total number of cases evaluated. It indicates the overall effectiveness of the model.

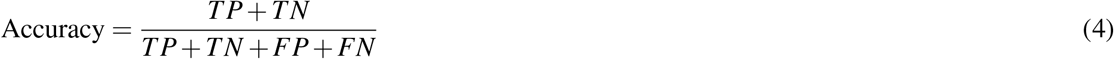
- **Cosine Similarity**: Measures the cosine of the angle between two vectors in an inner product space. It quantifies how similar the vectors are by determining if they point in approximately the same direction.

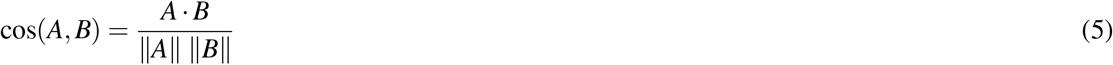

### *In silico* spectral database to validate annotation of empirical spectra

**Data** To build the *in silico* spectra database, we used the previously compiled list of SMILES of 30,591 unique compounds with associated experimental spectra. We also downloaded SMILES of metabolites from various online sources: Human Metabolome Database (HMDB) (217,760 SMILES), Chemical Entities of Biological Interest (ChEBI) (51,112 SMILES), ChEMBL (1,773,996 SMILES), Computational Toxicology (CompTox) (8,963 SMILES), NORMAN Suspect List Exchange (NORMAN-SLE) (114,051 SMILES), and NORMAN Substance Database (NORMAN SusDat) (106,632 SMILES). Additionally, we considered 42,648 SMILES from the NIST20, Agilent METLIN Metabolomics, MSDial, and GNPS databases. After removing duplicates, the initial database consisted of 2,211,691 unique SMILES.

To be as exhaustive as possible in the annotation of unidentified spectra from ARUS, we enlarged the *in silico* database with all compounds available for download from PubChem, resulting in an extended database of 96,492,904 predicted spectra.

To ensure consistency across the SMILES obtained from different databases, we calculated their canonical SMILES using the Rdkit package (https://www.rdkit.org). This process allowed us to identify how many SMILES corresponded to the same compound.

## Acknowledgements

This research was supported by projects PID2022-142600NB-I00 (MS-P and RGu), PID2022-136226OB-I00 (OY), and FPI grant PRE2020-095552 (OCT), from MCIN/AEI/10.13039/501100011033 and ERDF/EU, by projects 2021SGR-633 (MS-P. and RGu) and 2021SGR842 (OY) from the Government of Catalonia, and by the European Union NextGenerationEU/PRTR. MFK has received funding from the European Union’s Horizon 2020 research and innovation programme under the Marie Sklodowska-Curie grant agreement No. 945413 and from Universitat Rovira i Virgili.

## Code availability

A Python implementation of SingleFrag and trained SingleFrag models are available from https://github.com/MaribelPR/SingleFrag.

## Author contributions statement

MS-P and RGu designed the research. MP-R and RGu wrote code. MP-R performed experiments. MP-R, MFK, RGi and JMB collected and processed spectral data. SJ performed the manual elucidation of ARUS spectra. MP-R, OY, MS-P, and RGu analyzed and discussed results. MP-R, RGi, OY, MS-P, and RGu wrote the paper.

## References

1. Beger, R. et al. Metabolomics enables precision medicine: “A White Paper, Community Perspective”. Metabolomics 12, 149 (2016).

2. Qiu, S. et al. Small molecule metabolites: Discovery of biomarkers and therapeutic targets. Sig. Transduct. Target Ther. 8, 132 (2023).

3. Rafiq, T. et al. Nutritional metabolomics and the classification of dietary biomarker candidates: A critical review. Adv. Nutr. 12, 2333–2357 (2021).

4. Bauermeister, A., Mannochio-Russo, H., Costa-Lotufo, L. V., Jarmusch, A. K. & Dorrestein, P. C. Mass spectrometry-based metabolomics in microbiome investigations. Nat. Rev. Microbiol. 20, 143–160 (2022).

5. Viant, M. R. et al. Use cases, best practice and reporting standards for metabolomics in regulatory toxicology. Nat. Commun. 10, 3041 (2019).

6. Stein, S. Mass spectral reference libraries: An ever-expanding resource for chemical identification. Anal. Chem. 84, 7274–7282 (2012).

7. Giera, M., Yanes, O. & Siuzdak, G. Metabolite discovery: Biochemistry’s scientific driver. Cell Metab. 34, 21–34 (2022).

8. Vinaixa, M. et al. Mass spectral databases for LC/MS and GC/MS-based metabolomics: State of the field and future prospects. Trends Anal. Chem. 78, 23–35 (2016).

9. Frainay, C. et al. Mind the gap: Mapping mass spectral databases in genome-scale metabolic networks reveals poorly covered areas. Metabolites 8, 51 (2018).

10. Blaženović, I., Kind, T., Ji, J. & Fiehn, O. Software tools and approaches for compound identification of LC-MS/MS data in metabolomics. Metabolites 8, 31 (2018).

11. Dührkop, K. et al. SIRIUS 4: A rapid tool for turning tandem mass spectra into metabolite structure information. Nat. Methods 16, 299–302 (2019).

12. Aguilar-Mogas, A., Sales-Pardo, M., Navarro, M., Guimerà, R. & Yanes, O. iMet: A network-based computational tool to assist in the annotation of metabolites from tandem mass spectra. Anal. Chem. 89, 3474–3482 (2017).

13. Wang, Y., Kora, G., Bowen, B. P. & Pan, C. MIDAS: A database-searching algorithm for metabolite identification in metabolomics. Anal. Chem. 86, 9496–9503 (2014).

14. Ruttkies, C., Schymanski, E. L., Wolf, S., Hollender, J. & Neumann, S. MetFrag relaunched: Incorporating strategies beyond in silico fragmentation. 8, 3 (2016). J. Cheminform. 8, 3.

15. Ruttkies, C., Neumann, S. & Posch, S. Improving MetFrag with statistical learning of fragment annotations. BMC Bioinf. 20, 376 (2019).

16. Wang, F. et al. CFM-ID 4.0: More accurate ESI-MS/MS spectral prediction and compound identification. Anal. Chem. 93, 11692–11700 (2021).

17. Wei, J. N., Belanger, D., Adams, R. P. & Sculley, D. Rapid prediction of electron–ionization mass spectrometry using neural networks. ACS Centr. Sci. 5, 700–708 (2019).

18. Young, A., Wang, B. & Röst, H. MassFormer: Tandem mass spectrum prediction for small molecules using graph transformers (2023). 2111.04824.

19. Hong, Y. et al. 3DMolMS: prediction of tandem mass spectra from 3D molecular confo rmations. Bioinformatics 39, btad354 (2023).

20. Jaeger, S., Fulle, S. & Turk, S. Mol2vec: Unsupervised machine learning approach with chemical intuition. J. Chem. Inf. Model. 58, 27–35 (2018).

21. Zhou, J. et al. Graph neural networks: A review of methods and applications. AI Open 1, 57–81 (2020).

22. Atz, K., Grisoni, F. & Schneider, G. Geometric deep learning on molecular representations. Nat. Mach. Intell. 3, 1023–1032 (2021).

23. Jiang, D. et al. Could graph neural networks learn better molecular representation for drug discovery? A comparison study of descriptor-based and graph-based models. J. Cheminform. 13, 12 (2021).

24. Simón-Manso, Y. et al. Mass spectrometry fingerprints of small-molecule metabolites in biofluids: Building a spectral library of recurrent spectra for urine analysis. Anal. Chem. 91, 12021–12029 (2019).

25. Wishart, D. S. et al. HMDB 5.0: the Human Metabolome Database for 2022. Nucleic Acids Res. 50, D622–D631 (2021).

26. Horai, H. et al. MassBank: A public repository for sharing mass spectral data for life sciences. J. Mass Spectrom. 45, 703–714 (2010).

27. Elapavalore, A. et al. Adding open spectral data to massbank and pubchem using open source tools to support non-targeted exposomics of mixtures. Environ. Sci.: Process. Impacts 25, 1788–1801 (2023).

28. Sawada, Y. et al. Riken tandem mass spectral database (respect) for phytochemicals: A plant-specific ms/ms-based data resource and database. Phytochemistry 82, 38–45 (2012).

29. Xing, S., Shipei ad Shen, Xu, B., Li, X. & Huan, T. BUDDY: Molecular formula discovery via bottom-up MS/MS interrogation. Nat. Methods 20, 881–890 (2023).

30. Tanimoto, T. T. Elementary mathematical theory of classification and prediction. Int. Bus. Mach. Corp. (1958).

